# The evolution of neurosensation drives the gain and loss of phenotypic plasticity

**DOI:** 10.1101/2021.11.29.470389

**Authors:** Emily Y. Chen, Diane K. Adams

## Abstract

Phenotypic plasticity is widely regarded as important for enabling species resilience to environmental change and for species evolution. However, insight into the complex mechanisms by which phenotypic plasticity evolves in nature has been limited by our ability to reconstruct evolutionary histories of plasticity. By using part of the molecular mechanism, we were able to trace the evolution of pre-feeding phenotypic plasticity across the class Echinoidea and identify the origin of plasticity at the base of the regular urchins. The neurosensory foundation for plasticity was ancestral within the echinoids. However, coincident development of the plastic trait and the neurosensory system was not achieved until the regular urchins, likely due to pleiotropic effects and linkages between the two colocalized systems. Plasticity continues to evolve within the urchins with numerous instances of losses associated with loss of sensory capabilities and in one case loss of neurons, consistent with a cost associated with maintaining these capabilities. Thus, evidence was found for the neurosensory system providing opportunities and constraints to the evolution of phenotypic plasticity.

## Introduction

Phenotypic plasticity is one of the most common phenomena of the living world (Pigliucci, 2005). Plasticity allows an individual to produce different phenotypes (forms, functions, or behaviors) from the same genotype. This environmentally-induced phenotypic variation contributes the overall variation that serves as the material for natural selection, facilitates invasion of new habitats, and enables acclimatization to variable environments (Pfennig et al., 2010; Agrawal, 2001). In addition to contributing to species and trait evolution, plasticity is also a trait subject to evolutionary processes. Because phenotypic plasticity requires genetically encoded molecular and cellular machinery to sense and induce changes in phenotypes, the ability to be plastic or not is heritable and subject to selection pressure. Consistent with this, the rate and magnitude of the response to the environment – i.e., the shape of the reaction norms – can differ between genotypes (Murren et al., 2015) and can be experimentally evolved (Scheiner, 2002; Garland and Kelly, 2006).

There are constraints – costs and limits – to the evolution of phenotypic plasticity that prevent achieving the ideal phenotype for a given environment and may prevent a trait from being plastic at all (Murren et al., 2015; DeWitt, et al., 1998). Generally, processes that hinder trait evolution such as limited genetic variation and gene flow will also hinder the evolution of plasticity (Schlichting and Pigliucci, 1998). Pleiotropic effects in which a gene for one trait is linked to a gene for plasticity of another trait can also limit an evolutionary response. The molecular and cellular machinery (enzymes, signaling molecules, etc.) required to detect the environment, process information, and invoke a structural response have costs to the organism (Murren et al., 2015; DeWitt et al., 1998). If these costs are substantial relative to any adaptive advantage, plasticity may be selected against and subsequently lost.

The neurosensory machinery required to detect the environment is likely to be one of the main costs of plasticity and could also limit the evolution of plasticity (Snell-Rood, 2013). However, despite recent attention to the costs of plasticity, quantification of costs has been challenging and evidence for a significant cost is limited (Van Kleunen and Fischer, 2005; Van Buskirk and Steiner, 2009; Steiner and Van Buskirk, 2008; Auld et al., 2009). Interpopulation comparisons suggest that sensory capabilities can evolve over ecological timescales (Tsuji et al., 2011; Bay and Palumbi, 2014). Further, rapid radiations of sensory receptor genes (Nei et al., 2008; Nozawa et al., 2007; Raible et al., 2006) and plasticity in neural networks (Abbott and Nelson, 2000; Andersen, 2003) could reduce any potential limitation. Thus, changes to existing neurosensory infrastructure may present evolutionary opportunities.

A comparative approach that characterizes the natural evolution of plasticity across taxa would allow for testing these hypotheses regarding the costs, limits and opportunities for plasticity. For example, if neurosensory components are costly, then losses of phenotypic plasticity to be associated with losses or simplifications of the nerves or sensory receptor repertoire would be expected. However, it can be difficult to take the first step of tracing the evolution of plasticity across phylogenies due to ambiguity between loss of plasticity and an ancestral state before plasticity (i.e., plasticity has not yet evolved). This challenge can be surmounted when part or all of the mechanism of plasticity is known. Though plasticity itself may be lost, remnants of the mechanism are likely to remain due to diminished selection pressure. For example, if predator-induced plasticity is lost in a species or line of *Daphnia*, artificially-induced expression of juvenile hormone may still produce a phenotype that mimics the predator-induced form (Miyakawa et al., 2010; Dennis et al., 2014).

We take advantage of knowledge of part of the mechanism for phenotypic plasticity in sea urchin larvae. The feeding structure of many species of sea urchin vary with food concentration throughout larval development, including during the pre-feeding stage (Boidron-Metairon, 1998; Miner, 2007; Hart and Strathmann, 1994; Byrne et al., 2008; Sewell et al. 2004). When food is abundant, post-oral arm length is shorter. When food is scarce, post-oral arm length is longer. Plasticity during the pre-feeding stage must be sensory driven, since food is not yet ingested (Miner, 2007; Adams et al., 2011). Although the sensory receptor remains unknown, it has been established that sensation of food initiates a dopamine signal which is received by a dopamine type-2 receptor to inhibit post-oral arm elongation; this optimizes arm development and associated feeding potential relative to maternal lipid expenditure (Adams et al., 2011). There are distinct phylogenetic limits to when this phenotypic plasticity in arm elongation could have first evolved in echinoderms. While both Echinoidea (urchins and sand dollars) and Ophiuroidea (brittle stars) have a pluteus larval form with skeletal supports, morphological and molecular phylogenies support these as convergent forms that evolved independently (Williamson, 2003; McIntyre et al., 2014; Littlewood and Smith, 1995). Thus, it is likely that the plasticity of the pluteus feeding arms (including the skeletal elements), also evolved independently.

Here, the gains and losses of pre-feeding phenotypic plasticity were traced across the echinoids using not only the phenotypic outcome but also part of the underlying developmental signaling mechanism to identify the origin of pre-feeding plasticity. The evolutionary dynamics of pre-feeding plasticity were characterized in arm elongation to test the hypothesis that neurosensation of the environmental cue constrains the evolution plasticity.

## Results

### Genesis of Phenotypic Plasticity

We surveyed echinoids for the pre-feeding response to food to determine when phenotypic plasticity evolved. Molecular and morphological data place cidaroids as the most basal extant taxa within Echinoidea (Littlewood and Smith, 1995; Kroh and Smith, 2010; Smith et al., 2006). We did not find evidence of shortened post-oral arms in the presence of food in the cidaroid *Eucidaris tribuloides* (Figure 1, Table 1). The lack of pre-feeding plasticity in *E. tribuloides* is not unexpected, due to a temporal mismatch between the timing of arm elongation and the onset of feeding. *E. tribuloides* begins feeding before the post-oral arms have substantially elongated. These results support a more recent origin of plasticity within the Echinoids.

**Figure 1:**
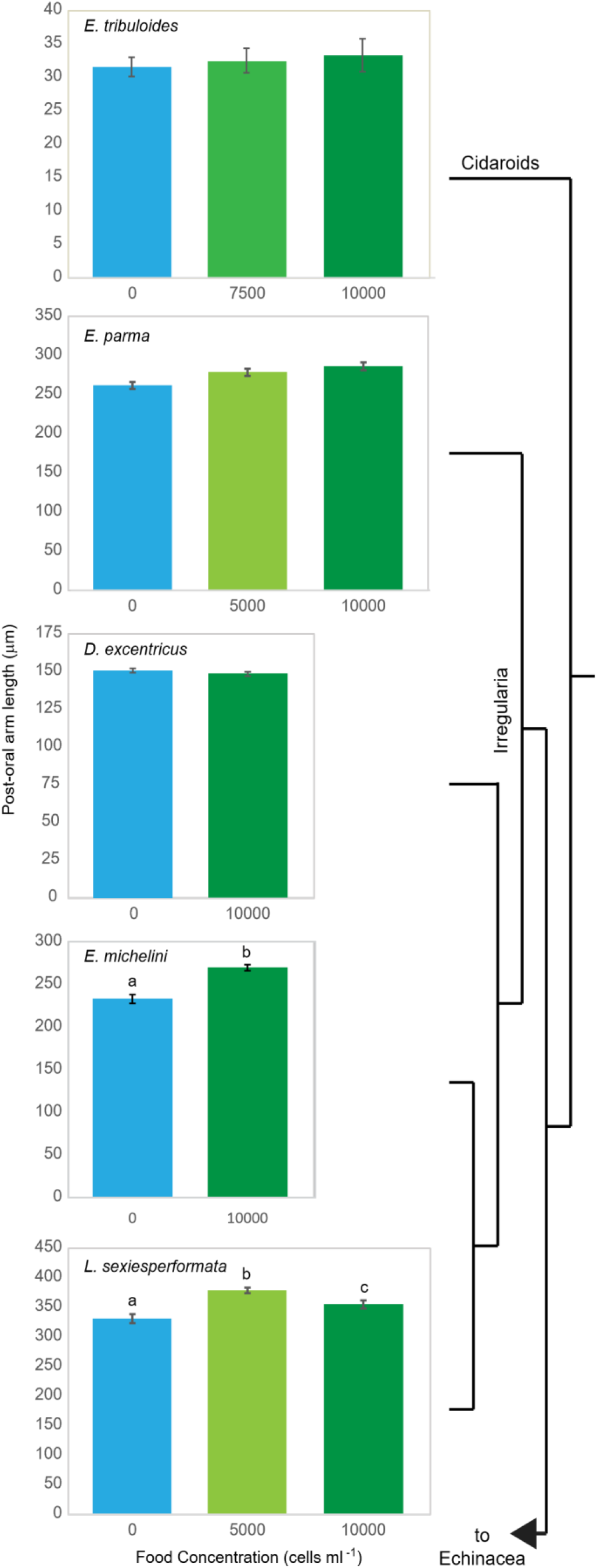
Canonical pre-feeding plasticity is absent in the basal Cidaroids and irregular urchins. Change in post-oral arm length at initiation of feeding averaged across families with food concentration in the Cidaroids and Irregularia. Phylogenetic tree is not scaled to divergence. Error bars, ± standard error of the mean. Letters denote a significant difference between food treatments at p < 0.05 (Table 1).

**Table 1:** Two-factor ANCOVAs with body rod (BR) as a covariate for irregular urchins.

|  | *E. parma* |  | *D. excentricus* |  | *L. sexiesperforata* |  |
| --- | --- | --- | --- | --- | --- | --- |
| Source | F statistic | p value | F statistic | p value | F statistic | p value |
| Family | $F_{1,213} = 0.250$ | 0.618 | **$F_{1,115} = 8.663$** | **0.004** | $F_{1,125} = 1.757$ | 0.187 |
| Food | $F_{2,213} = 1.434$ | 0.241 | $F_{1,115} = 1.010$ | 0.316 | **$F_{2,125} = 15.697$** | **0.000** |
| Family x Food | **$F_{2,213} = 7.561$** | **0.001** | **$F_{1,115} = 5.895$** | **0.016** | **$F_{2,125} = 4.107$** | **0.019** |
| BR | $F_{1,213} = 1.695$ | 0.194 | $F_{1,115} = 0.069$ | 0.793 | **$F_{1,125} = 24.065$** | **0.000** |

Irregular urchins have elongated arms during the pre-feeding stage and some are known to alter the length of their post-oral arms in response to food after feeding starts (Boidron-Metairon, 1988; Reitzel and Heyland, 2007). However, we did not observe the canonical plastic response in any of the irregular species tested, *Echinarachnius parma*, *Dendraster excentricus*, *Encope michelini*, and *Leodia sexiesperforata* (Figure 1). Post-oral arm lengths were significantly different in the presence of food for the keyhole sand dollars *E. michelini* (Student’s t-test, p < 0.001) and *L. sexiesperforata* (F_2,125_ = 15.697, p < 0.001) (Figure 1, Table 1). However, the response was in the opposite direction of the previously described canonical response. Those larvae exposed to abundant food had significantly longer post-oral arms than those without food. This elongation may be a specific response of the keyhole sand dollars (Mellitidae), although changes in post-oral arm length in response to food concentration followed a similar, but non-significant trend for *E. parma* (F_2,213_ = 1.434, p = 0.241, Table 1).

Surprisingly, pre-feeding plasticity was not detected for the common sand dollar *D. excentricus* in our experiments (Fig 1, F_1,115_ = 1.010, p = 0.316). *D. excentricus* has demonstrable phenotypic plasticity after feeding starts (Boidron-Metairon, 1988; Hart and Strathmann, 1994) and has been previously reported to have the canonical pre-feeding response (Miner, 2007). The differences between our observations and those of Miner, (2007) could be due to the different populations tested (Goleta, CA vs Orcas Island, WA) or our ability to detect the small magnitude of change (~5 % reduction). However, the lack of the canonical plasticity in *D. excentricus* is consistent with the results for the other irregular urchins.

Significant pre-feeding phenotypic responses to food abundance were only detected within the regular urchins. *Arbacia punctulata, Lytechinus variegatus*, and *Strongylocentrotus purpuratus* all had significantly shorter post-oral arm lengths in the presence of high food (Figure 2, Table 2). Three other taxa tested, *Echinometra lucunter, Lytechinus pictus*, and *Lytechinus variegatus carolinus*, did not significantly respond to changes in food concentration (Figure 2, Table 2). The basal position of *A. punctulata* within the regular urchins supports the interpretation that plasticity is ancestral within the clade and that there have been multiple losses. However, the alternative of multiple convergent evolutionary events within the regular urchins is also a possibility based on these data.

**Figure 2:**
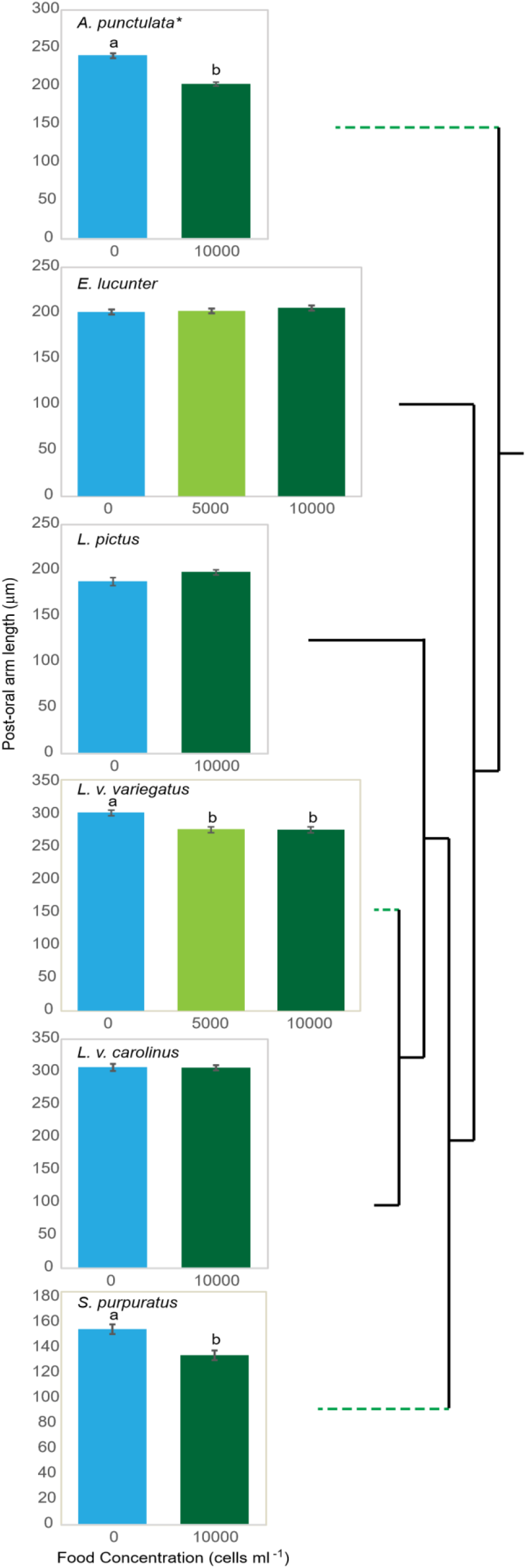
Pre-feeding plasticity has dynamically evolved within the regular urchins. Change in post-oral arm length at initiation of feeding averaged across families with food concentration in Echinacea, the regular urchins. *Data from a single family of *A. punctulata* is presented for clarity, though food treatment was significant across all families tested (Table 2). Phylogenetic tree is not scaled to divergence. Green dotted lines denote taxa with canonical pre-feeding plasticity. Error bars, ± standard error of the mean. Letters denote a significant difference between food treatments at p < 0.05 (Table 2).

**Table 2:** Two-factor ANOVAs with body rod (BR) as a covariate for regular urchins.

|  | *A. punctulata* |  | *E. lucunter* |  | *L. pictus* |  |
| --- | --- | --- | --- | --- | --- | --- |
| Source | F statistic | p value | F statistic | p value | F statistic | p value |
| Family | **$F_{3,336} = 163.095$** | **0.000** | **$F_{1,345} = 16.847$** | **0.000** | $F_{2,86} = 0.320$ | 0.727 |
| Food | **$F_{1,336} = 6.003$** | **0.015** | $F_{2,345} = 1.046$ | 0.352 | $F_{1,86} = 3.664$ | 0.059 |
| Family x Food | **$F_{3,336} = 6.734$** | **0.000** | $F_{2,345} = 2.103$ | 0.124 | $F_{2,86} = 0.223$ | 0.801 |
| BR | **$F_{1,336} = 26.944$** | **0.000** | **$F_{1,345} = 8.628$** | **0.004** | $F_{1,86} = 0.610$ | 0.437 |

|  | *L. v. variegatus* |  | *L. v. carolinus* |  |
| --- | --- | --- | --- | --- |
| Source | F statistic | p value | F statistic | p value |
| Family | **$F_{1,256} = 8.837$** | **0.003** | **$F_{1,67} = 19.445$** | **0.000** |
| Food | **$F_{2,256} = 14.362$** | **0.000** | $F_{1,67} = 0.019$ | 0.892 |
| Family x Food | **$F_{2,256} = 21.356$** | **0.000** | $F_{1,67} = 0.002$ | 0.968 |
| BR | **$F_{1,256} = 13.757$** | **0.000** | $F_{1,67} = 0.512$ | 0.477 |

### Remnant Signaling Mechanism

To distinguish between evolutionary losses and convergent gains, *E. lucunter, L. pictus*, and *L. variegatus carolinus* were tested to see if they still retained a phenotypic response to activation of dopamine type-D_2_ receptors (DRD2) even though they had lost the response to food. Dopamine signaling through DRD2 is required for the presence of food to inhibit arm elongation in the regular urchin *S. purpuratus* (Adams et al., 2011). If plasticity is ancestral within the regular urchins, we would expect that all of the regular urchins would use this same neural signaling mechanism and that even those that lost the plastic response might still retain this signaling remnant. Alternatively, if plasticity evolved convergently multiple times, differences are expected in the neural signaling mechanism and no response to dopamine signaling in those species without plasticity. Consistent with an ancestral origin within the regular urchins, activation of dopamine type-D_2_ receptors with the selective agonist, quinpirole, inhibited post-oral arm elongation in all of the regular urchins tested, including those without the response to food (Figure 3).

**Figure 3.**
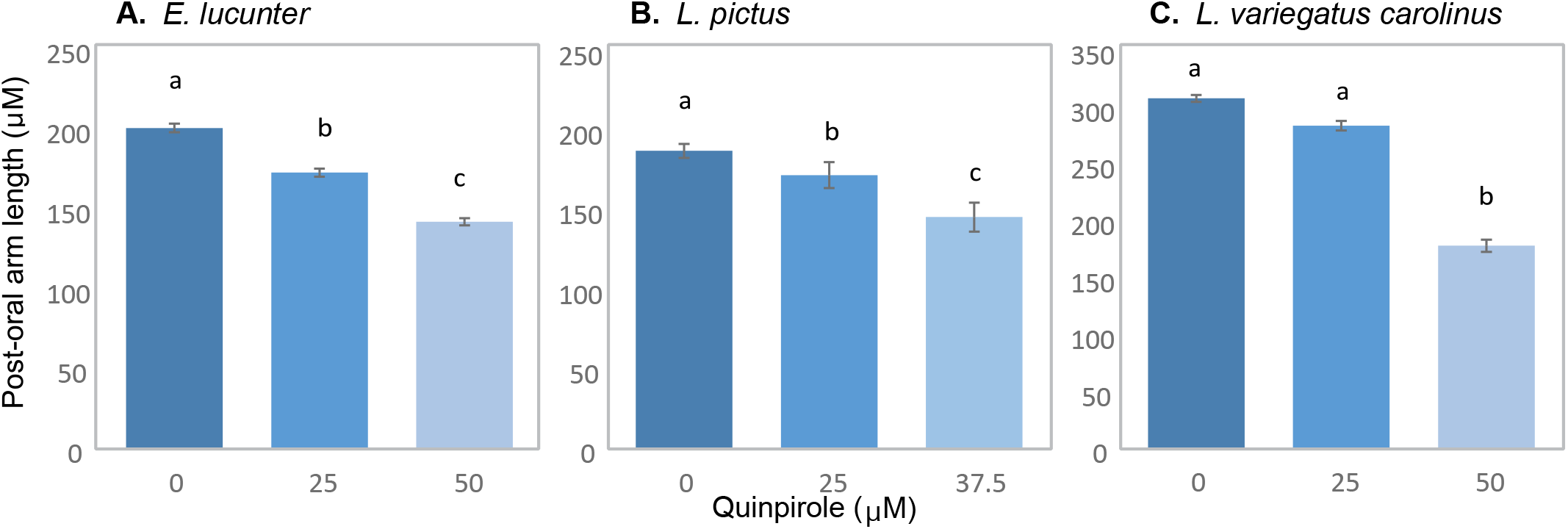
Regular urchins without phenotypic plasticity retain the phenotypic response to dopamine receptor activation. Change in post-oral arm length at initiation of feeding with treatment of the dopamine type-2 receptor agonist, Quinpirole, at varying concentrations for the three regular urchins, *E. lucunter* (A; F2,348 = 124.996, p < 0.001), *L. pictus* (B; F2_,96_ = 38.662, p < 0.001), and *L. variegatus carolinus* (C; F2,100 = 290.433, p <0.001) lacking a phenotypic response to food (Figure 2). Error bars, ± standard error of the mean. Letters denote significant post-hoc Bonferroni comparisons between Quinpirole treatments, p < 0.05.

### Dopaminergic neural development

To test whether neural development constrained the evolution of phenotypic plasticity, we characterized the temporal and spatial development of putative dopaminergic neurons (TH-positive) throughout Echinoidea. The member of the most basal group, *E. tribuloides*, developed TH-positive lateral ganglia near the future post-oral arms before feeding and before arm elongation (Figure 4A). After feeding starts, TH-positive neurons are also detected in the oral ganglia around the mouth and associated with the stomach. Thus, the requisite neural developmental systems were in place ancestrally, before the evolution of pre-feeding plasticity.

**Figure 4:**
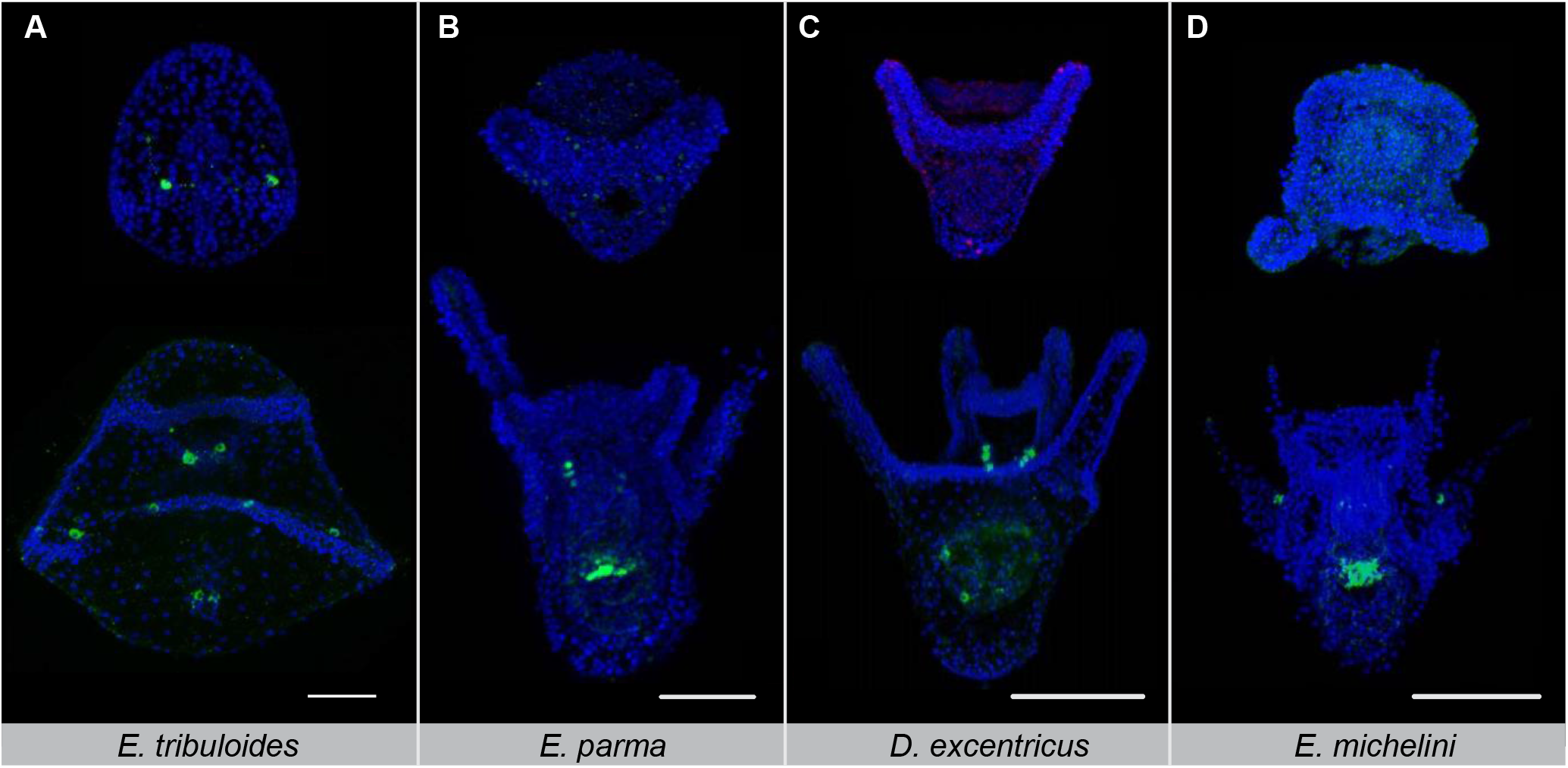
Dopaminergic development in the lateral ganglia was ancestrally present then lost. Immunodetection of the dopamine biosynthesis enzyme tyrosine hydroxylase (green) at prism or early pluteus stage (top row) and after feeding starts (bottom row) for the cidaroid, E. tribuloides (A), and irregular urchins, E. parma (B), D. excentricus (C), and E. michelini (D). DAPI counterstain, Blue. Scale bar, 100 μm for all images.

However, the irregular urchins investigated have altered dopaminergic development (Figure 4B-D). Tyrosine hydroxylase first appears after feeding begins in *D. excentricus*, *E. parma*, and *E. michelini*. In both *D. excentricus and E. parma*, TH-positive neurons are detected in the mouth and gut, but not as lateral ganglia near the post-oral arms. Since we can detect TH-positive cells in the mouth and gut, we do not believe that the absence of TH-positive lateral ganglia is due to a detection issue. The lack of early dopaminergic lateral ganglia is consistent with the lack of feeding arm plasticity detected within the irregular urchins (Figure 1) and suggests that neural development may have constrained the evolution of pre-feeding plasticity within this clade.

Only within regular urchins does the development of the post-oral arms and TH-positive neurons coincide during the pre-feeding stage. Lateral dopaminergic neurons developed during the prism stage, at approximately the time of arm elongation in *A. punctulata, L. pictus, L. variegatus, L. variegatus carolinus*, and *S. purpuratus* (Figure 5). At the onset of feeding, TH-positive neurons appear around the mouth as oral ganglia and begin to appear in the stomach. The number of TH-positive neurons associated with lateral ganglia near the post oral arms, vary between species. Both *A. punctulata* and *L. variegatus* subspp. develop multiple TH-positive neurons along the post-oral arms. Fewer TH-positive neurons develop in *L. pictus* and the shorter *S. purpuratus* arms. *Echinometra lucunter* is the exception – this species does not develop TH-positive neurons until post-feeding and even then, the lateral ganglia appear to be absent. This change in neural development may be responsible for the loss in pre-feeding phenotypic plasticity in *E. lucunter*. Thus, multiple distinct changes in neural development – timing (heterochrony), number (heterometry), and location (heterotopy) – could have contributed to the evolutionary constraints and opportunities for pre-feeding phenotypic plasticity in sea urchin larvae.

**Figure 5:**
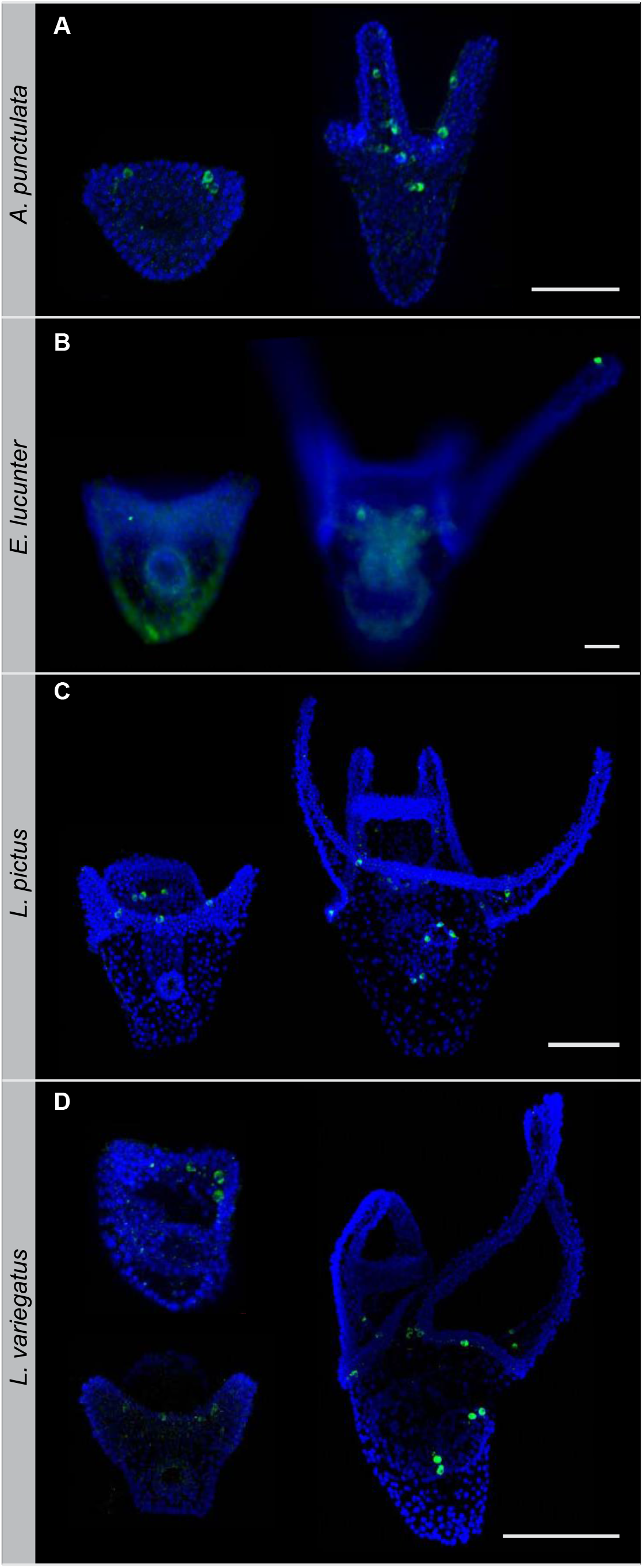
Dopamine neurons develop in the lateral ganglia occurs in most regular urchins. Immunodetection of the dopamine biosynthesis enzyme tyrosine hydroxylase (green) at prism or early pluteus stage (left) and after feeding starts (right) for the regular urchins, *A. punctulata* (A), *E. lucunter* (B), *L. pictus* (C), and the *L. variegatus* subsp. (D). The prism stage is shown for *L. variegatus carolinus* (top) and *L. variegatus variegatus* (bottom). DAPI counterstain, Blue. Scale bar, 100 μm all images.

## Discussion

### Origin of Pre-feeding Phenotypic Plasticity

The data suggest that pre-feeding phenotypic plasticity of the post-oral arms arose in the regular urchins and has continued to evolve within the clade. While there were significant responses to food for some irregular urchins, the response was in the opposite direction of the established response. Consistent with this departure from the plastic response observed in regular urchins, all of the irregular urchins lacked putative dopaminergic neurons in the lateral ganglia during the pre-feeding stage. This includes *Dendraster excentricus* which was previous reported to have a subtle pre-feeding response to food (Miner, 2007). However, *D. excentricus* and *S. purpuratus* responded morphologically to different cues (soluble vs algal bound, respectively), which is consistent with convergent evolution (Miner, 2007).

An evolutionary origin of pre-feeding plasticity at the base of the regular urchins is in contrast to phenotypic plasticity that occurs after feeding starts, when additional and more reliable cues, such as metabolic byproducts, could be used to assess food availability. Feeding plasticity has not been reported for any of the basal echinoids tested to date (2 of 2 cidaroids (McAlister, 2008) and 3 of 3 diademids (McAlister, 2008; Soars et al., 2009)). However, both the irregular (3 of 5 species (Boidron-Metairon, 1988; Hart and Strathmann, 1994; Reitzel and Heyland, 2007; Eckert, 1995)) and regular (9 of 12 species (Miner and Vonesh, 2004; Strathmann et al., 1992; Poorbagher et al., 2010; Bertram and Strathmann, 1998; Miner, 2005; McAlister, 2007)) urchins have taxa that exhibit phenotypic responses to food after feeding starts. Interestingly, the two species of irregular urchins reported to lack feeding plasticity are the mellitid keyhole sand dollars *E. michelini*, and *L. sexiesperforata*, which also lacked canonical pre-feeding plasticity here (Reitzel and Heyland, 2007). Similarly, the three species of regular urchins lacking post-feeding plasticity were species in the genus *Echinometra*, including *E. lucunter*, which also lacked canonical pre-feeding plasticity here (McAlister, 2008). This may be a recent loss isolated to the genus *Echinometra*, as feeding plasticity was reported in the Echinometrid *Heliocidaris tuberculate* (Soars et al, 2009). Knowledge of the mechanism(s) underlying plasticity during the planktonic feeding stage would again provide the ability to discriminate between evolutionary losses and multiple convergent gains.

### Evolutionary Opportunity and Constraint by Neural Systems

The results suggest that development of the dopaminergic neurons in the lateral ganglia was already in place within the ancestral echinoids, the cidaroids. This provides a foundational component that could have later facilitated the evolution of pre-feeding phenotypic plasticity. However, in the cidaroids, post-oral arm elongation does not occur until days after feeding starts, so the timing of skeletal elongation may have been an ancestral constraint on the evolution of pre-feeding plasticity.

We propose that pleiotropic effects or gene linkage associated with the temporal shift in arm elongation, altered development of the dopaminergic neurons in the lateral ganglia (Figure 6). This would explain the loss or temporal shift in development of dopaminergic lateral ganglia in the irregular urchins investigated. The lateral ganglia develops within the lateral/boundary ectoderm, where epithelial-mesenchymal signaling is known to coordinate skeletal elongation (McIntyre et al., 2013, 2014; Adomako-Ankomah and Ettensohn, 2013; Duloquin et al., 2007; Ettensohn, 2009). Thus, it is possible that changes in the signaling milieu to advance skeletal elongation could have suppressed dopaminergic development. In support of this, many of the genes within the skeletogenic gene regulatory network (Rafiq, 2014), including FGF, Pax 2/5/8, Wnt5, and Otp (McIntyre et al., 2013; Röttinger et al., 2008; Cavalieri et al., 2003), also have roles in dopaminergic development in other systems (Hegarty et al., 2013; Ryu et al., 2007; Smidt et al., 2003). A decoupling of gene expression or function in the regular urchins would be necessary to allow for the coincident development of dopaminergic lateral ganglia and the post-oral arms during the pre-feeding stage. The loss of plasticity in the regular urchin *E. lucunter* could represent a reversion to the irregular-like state with early skeletal elongation and delayed neural development. Thus, dynamic changes in the development of the lateral ganglia throughout echinoidea are likely to have both constrained and provided opportunity for plasticity.

**Figure 6.**
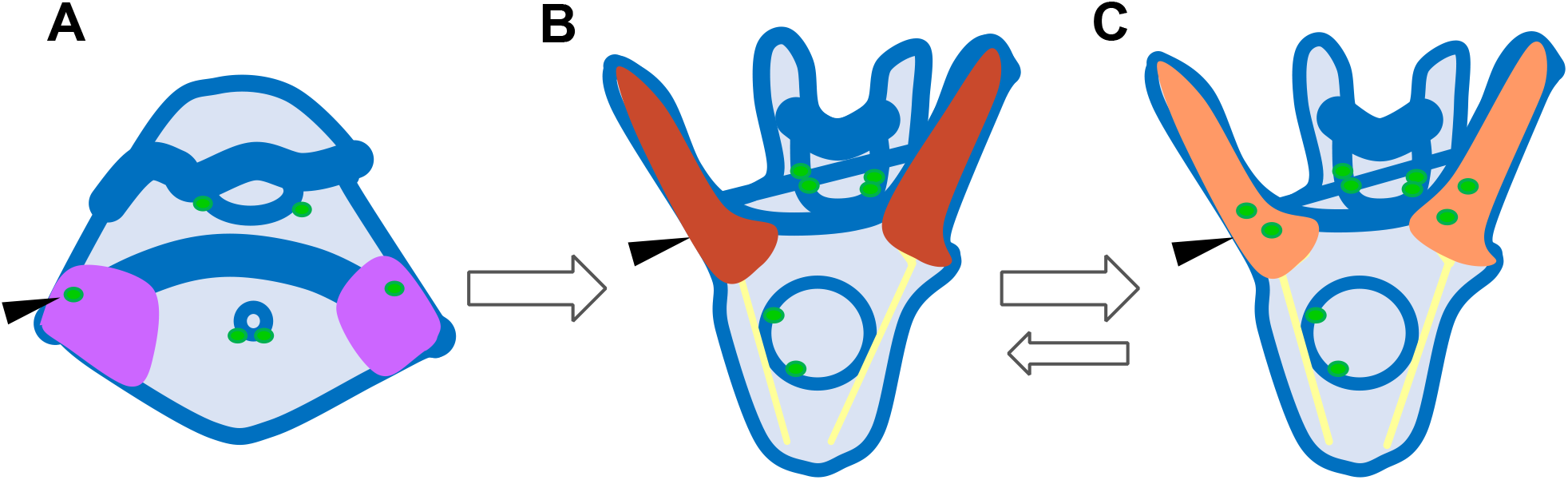
Model for the evolution of post-oral arm elongation and dopaminergic development. The change in the signaling milieu [pink (A) to red (B)] that allowed for earlier elongation of the post-oral arms likely also inhibited the early development of dopaminergic neurons (green circles) in the lateral ganglia (black arrows). Another shift in signaling or relaxation of pleiotropy at the base of the regular urchins restored early dopaminergic development (C). Dynamic evolution within the regular urchins suggests that there may also be shifts back to the prior evolutionary state (B).

### Evolution of Neurosensation

Changes in neural development cannot account for the differences in plasticity within all of the regular urchins. *L. variegatus carolinus* and *L. pictus* develop TH-positive cells at the appropriate time and place (Figure 5 B, C) and respond to activation of DRD2 (Figure 3 B, C). However, they lack the pre-feeding phenotypic response to food concentration (Figure 2). This suggests that the change responsible for the loss of plasticity occurred upstream of the dopamine receptor – during the neurosensory process.

Given that there is evidence for rapid evolution of putative sensory receptors in sea urchins (Raible et al. 2006) that could affect developmental plasticity, we hypothesize that changes in sensory receptor expression or sequence caused the loss of arm plasticity. Sea urchins have a large repertoire of GPCRs and immune receptors that could act in the sensation of food (Rast et al., 2006; Hibino et al., 2006). Both immunity receptors and GPCRs are often found in large tandem arrays of genes and pseudogenes suggestive of gene duplications. In the purple sea urchin, 979 GPCRs have been identified – comprising nearly 3% of the predicted proteins. Two groups of these GPCRs have rapidly expanded and are most similar to vertebrate olfactory receptors (Raible et al., 2006). Although innate immunity receptors are generally believed to have ancient origins and minimal subsequent evolution, there is genomic evidence from sea urchins for extensive radiations in this group as well, with 10-20 fold more genes in the purple sea urchin than in humans (Rast et al., 2006; Hibino et al., 2006). Rapid evolution of sensory receptors is also consistent with the recent evolutionary loss of the response between relatively close sister species (~3 million years) and subspecies (less than a million years) in the genus *Lytechinus* (Zigler and Lessios, 2004). However, the identity of the sensory receptor and its evolution remains to be determined.

## Conclusion

The data demonstrate the power of a comparative approach to understand the evolutionary dynamics of phenotypic plasticity when part of the molecular mechanism is known. Once within an evolutionary context, we were able to assess the role of neural development in constraining the evolution of plasticity. In this case, ancestral neural development provided a foundational opportunity, rather than constraint. Instead, we propose that interactions between neural development and development of the plastic trait constrained the rise of phenotypic plasticity and decoupling was necessary to allow for the advent of plasticity. Once established, phenotypic plasticity has continued to evolve dynamically both through changes in neural development and potentially evolution of sensory receptors.

## Materials & Methods

### Embryo and larval culture

Adult echinoids were obtained from the following vendors for broodstock: *Lytechinus variegatus* (Tom’s Caribbean and Reeftopia, Florida Keys, FL), *L. variegatus carolinus* (Duke Marine Labs, Beaufort, NC), *L. pictus* (Marinus, Goleta, CA), *Echinometra lucunter* (Reeftopia, Florida Keys, FL), *Arbacia punctulata* (Gulf Specimen Marine Lab, Panacea, FL and Duke Marine Lab, Beaufort, NC), *Dendraster excentricus* (Marinus Scientific, Goleta, CA), *Echinarachnius parma* (MBL, Woods Hole, MA), *Encope michelini* (Reeftopia, Florida Keys, FL), *Leodia sexiesperformata* (Reeftopia, Florida Keys, FL), and *Eucidaris tribuloides* (Tom’s Caribbean and Reeftopia, Florida Keys, FL). Gametes were obtained using intracoelomic injections of 0.55 M KCl. Embryos were cultured using standard methods at densities of 1-5 embryos ml^−1^ in artificial seawater (ASW) at 21 °C for tropical species or 15 °C for temperate species. Larvae were treated with 5,000, 7,500 or 10,000 cells ml^−1^ of the algae *Dunaliella* sp. to assay for the developmental-response to food. Algal concentration was determined using a hemocytometer. Larvae of the regular echinoids were also treated with the specific type-D2 receptor agonist (Maggio and Millan, 2010), quinpirole, at late gastrula stage or prism stage to test for conservation of the dopamine-signaling mechanism. Doses of 0, 25, and 50 μM were used. The highest dose was decreased to 37.5 μM for *L. pictus* due to sickness in this species at 50 μM.

### Quantification of skeletal lengths

Post-oral arm and body rod lengths were assayed just before feeding begins as in Adams et al., (2011). The time post fertilization varied with each species and was experimentally determined by observing algal particles within the gut. All collections were done when algae were observed in less than 50% of the larvae’s guts. Larvae were randomly sampled from each treatment, such that sample sizes varied but all were n ≥ 20 individuals each. Larvae were squash mounted on microscope slides to position the skeletal elements in the same plane, then imaged on a Zeiss Axiovert 200M or Zeiss Axiovert A1 inverted microscope at 20x under differential inference contrast (DIC) which readily identifies the birefringent skeletal elements. The skeletal lengths were quantified from the digital images using Zen Lite software (Carl Zeiss MicroImaging).

### Statistical Analyses

The response of post-oral arm length to algal and quinpirole treatments was assessed using a two-way ANCOVA, where perturbation treatment (food or quinpirole) and biological replicate (male-female cross) were fixed effects. Body rod length was included as a covariate. We used post-hoc Bonferroni-corrected pair-wise comparisons when effects were significant at p < 0.05. Experiments were replicated with two or more sets of non-related full siblings (male-female crosses) for all species except *E. michelini* and *L. sexiesperformata*, due to limitations in obtaining ripe broodstock. For these species, only one male and female were available yielding one set of full siblings; thus, a one-way ANOVA (*E. michelini*) or Student’s two-tailed t-tests (*L. sexiesperformata*) were used. All datasets were determined to be normal based on probability distribution plots. All statistical analyses were done in SYSTAT v10 with output to three decimal places, thus exact *P* values are given if *P* > 0.001.

### Immunofluorescent staining

Immunostains for tyrosine hydroxylase (1:200, ImmunoStar #22941) were performed as in Adams et al., (2011) on two stages of larvae: 1) just after the initial elongation of the post-oral arms and 2) after feeding started. When tyrosine hydroxylase was not detected at these developmental stages, later stage larvae were also assayed to ensure that the antibody worked in all species tested. Specificity of the antibody in echinoids was established in *S. purpuratus* by morpholino knock down of tyrosine hydroxylase. Larvae were imaged using a Zeiss Axiovert 200M epifluorescent inverted microscope with an optically sectioning ApoTome unit or Zeiss LSM 710 Confocal microscope at 20x or 40x. Stacked images were prepared using Imaris (Bitplane Inc., St. Paul, MN).

## Acknowledgements

We are grateful for discussions with R. Range and A. Sethi. Support for this research was provided by Rutgers, the State University of New Jersey.

## Authors Contributions

E.Y.C. led the execution of the experiments and data analysis, and contributed to manuscript preparation. D.K.A. led the design of the experiments, supervised execution of the experiments and data analysis, and contributed to manuscript preparation.

